# Broad compatibility between yeast UAS elements and core promoters and identification of promoter elements that determine cofactor specificity

**DOI:** 10.1101/2022.11.03.515066

**Authors:** Jeremy A. Schofield, Steven Hahn

## Abstract

Three general classes of yeast protein-coding genes are distinguished by their dependence on the transcription cofactors TFIID, SAGA and Mediator (MED) Tail, but little is known about whether this dependence is determined by the core promoter, Upstream activation sites (UASs), or other gene features. It is also unclear whether UASs can broadly activate transcription from the different promoter classes or whether efficient transcription requires matching UASs and promoters of similar gene class. Here we measure transcription and cofactor specificity for tens of thousands of UAS-core promoter combinations. We find that <5% of UASs display strong core promoter specificity while most UASs can broadly activate promoters regardless of regulatory class. However, we find that matching UASs and promoters from the same gene class is generally important for optimal expression. From examining the cofactor dependence of this large UAS-promoter set, we find that sensitivity to rapid depletion of MED Tail or SAGA is dependent on the identity of both UAS and promoter while dependence on TFIID localizes to only the core promoter. Our results explain why transcription factor-mediated MED recruitment to the UAS does not always result in Tail-dependent transcription and highlight the role of TATA and TATA-like promoter sequences in MED Tail function.

## Introduction

Regulation of transcription is vital for establishing organismal complexity and in mounting responses to cellular stresses. Eukaryotic genes display distinct transcriptional behaviors dependent on regulatory classes. For example, inducible (or developmentally regulated) metazoan genes tend to be transcribed with high variance and are enriched for TATA boxes in their core promoter, whereas constitutive (or housekeeping) genes are transcribed at lower levels, typically with low variance^1–4^. Understanding the differences in regulatory mechanisms utilized at these different gene classes is important but the molecular basis of class-specific gene regulation remains uncertain.

Transcriptional cofactors (coactivators, corepressors and chromatin modifying factors) function as intermediates between transcription factors that bind at enhancers and the transcription machinery^5–13^. Cofactor targets range from chromatin to basal transcription factors and ultimately regulate formation of the pre-initiation complex (PIC) at the core promoter as well as post initiation events^14,15^. Recent results in yeast, using rapid cofactor depletion and nascent transcriptome analysis, refined the gene-specific targets of several cofactors and identified three fundamental classes of yeast genes^16–18^. Transcription of most yeast genes (~87% of yeast protein coding genes) is highly dependent on TFIID and the bromodomain-containing factors Bdf1/2 but is largely insensitive to rapid depletion of SAGA and the activator binding domain of Mediator (MED) termed Tail. These TFIID-class genes show predominantly lower expression and are depleted for TATA boxes in their promoters, analogous to metazoan housekeeping genes. Expression from another subset of yeast genes, (~13%) termed coactivator redundant (CR, analogous to developmentally regulated genes), is moderately sensitive to rapid depletion of either SAGA or TFIID and largely insensitive to Bdf1/2 depletion. However, simultaneous depletion of both TFIID and SAGA causes a severe decrease in transcription of these genes. About one half of the CR genes (~6% of total genes) are also sensitive to rapid MED Tail depletion (Tail-dependent genes)^18^. The differential cofactor sensitivity of all these genes likely reflects different pathways used in formation of the PIC. Whether these yeast gene classes (TFIID, CR/Tail-dependent, CR/Tail-independent) are defined by the function of their enhancer/UAS elements, core promoters, or a combination of the two is an important open question.

Seminal work established that enhancers can display specificity in activation of core promoters containing specific regulatory motifs.^19,20^ Several later reports used massively parallel reporter assays (MPRAs, eg. STARR-seq^21^) to investigate whether enhancer-core promoter specificity is a general phenomenon in metazoans. Work in the *Drosophila* system suggested that many enhancers display high functional specificity for promoters with optimum expression requiring paired elements from either the developmental or housekeeping gene classes^22^. In contrast, recent work in human and mouse systems suggests that most enhancers can activate a wide range of core promoters and therefore display low specificity^23,24^. These later studies found that selectivity in enhancer activity at various core promoters is subtle and largely corresponds to matched enhancers and promoters from the same regulatory class. In addition, the function of some enhancers display specificity for different cofactors^25^, but the mechanistic basis of the functional interplay between enhancers, promoters, and coactivators remains poorly defined.

Metazoan enhancers can be megabases distant from the core promoters they regulate^26^, and the spatial relationship of enhancers and promoters is an additional layer of complexity^27,28^. The yeast genome is an ideal context to study the functional connection between distal and core promoter elements, as long-distance regulation of core promoters by UAS elements, the sites for the binding of sequence specific transcription factors, has not been described^29^. However, this arrangement presents a challenge in the design of a scalable assay to study the transcription of many regulatory sequences since UAS elements placed downstream of a core promoter are unlikely to self-transcribe and therefore be identified in a traditional STARR-seq assay. Here we developed a large-scale reporter assay measuring the transcriptional activities and coactivator sensitivity for tens of thousands of yeast UAS sequences and core promoter combinations. Our results are consistent with a model in which most UASs can broadly activate different core promoter classes but display quantitative specificity in expression level that depends on matching UASs and promoters of similar class. In contrast, we find that only a small fraction of UASs (~5 %) show strict specificity for activating the TFIID or CR class genes. Finally, we find that SAGA and MED-Tail specificity appear dependent on appropriate matching of both the UAS and core promoter, whereas TFIID specificity appears primarily dependent on the core promoter alone.

## Results

### A reporter assay to assess functional matching of yeast UAS and core promoter regions

To test whether yeast UASs preferentially activate transcription from different classes of core promoters, we devised a large-scale combinatorial cloning strategy to generate tens of thousands of UAS-core combinations driving expression of a barcoded reporter gene. Four core promoters from three separate gene classes (based on SAGA, TFIID, and MED-Tail sensitivity^16,18^) were cloned in combination with a majority of all yeast UASs (~4200) or random genomic controls (**Fig 1A, S1A-B**, see Methods for cloning and region definitions). The *KRS1* and *SIT1* genes are both TATA-containing CR class genes but only the endogenous *SIT1* gene is MED Tail-dependent. Both *NOP13* and *RPC10* are GA element (GAE)- containing TFIID class genes. The GAE (sequence GAAAA) is found at nearly half of TATA-less promoters in *S. cerevisiae* and was shown to be important for TBP binding and promoter function^30^. The UAS-core libraries were transformed into yeast harboring the degron-tagged coactivator subunits Spt7 (SAGA), Taf13 (TFIID), or Med15 (MED-Tail), and we assessed both baseline transcription for each UAS-core combination and changes in transcription following rapid coactivator depletion (**Fig 1B**). Expression levels for each UAS-core combination was determined by normalizing RNA barcode signal by corresponding plasmid DNA signal from transformed yeast, and expression levels are highly reproducible between replicates (see Methods).

**Fig1.**
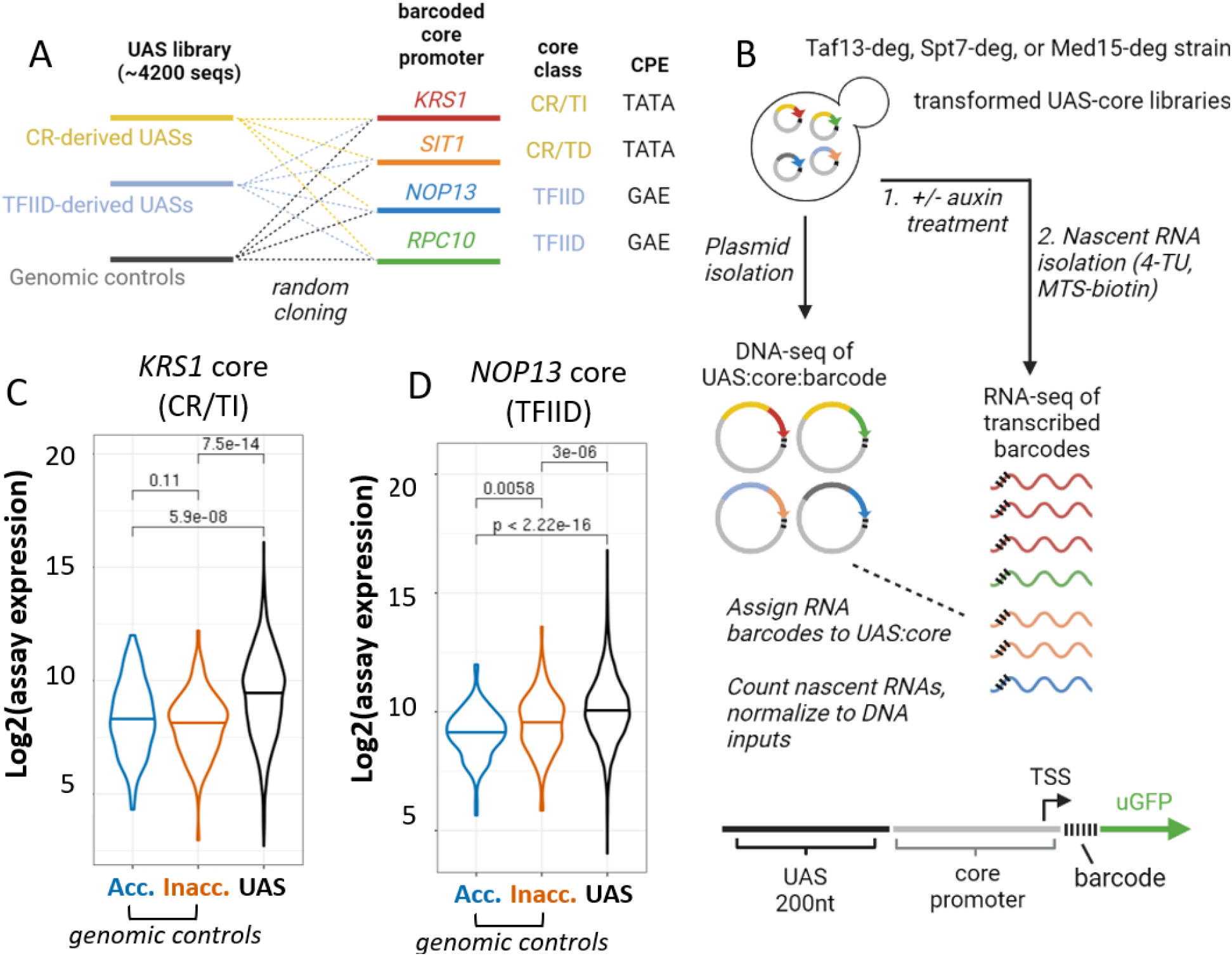
A large scale transcriptional reporter assay to measure expression and coactivator sensitivity for tens of thousands of UAS-core promoter combinations. **A)** Yeast genomic UASs (200nt) and random genomic sequences are cloned upstream of four core promoters from varying coactivator classes (Coactivator Redundant class, CR; TFIID-class, TFIID; Mediator Tail-independent, TI; Mediator Tail-dependent, TD). Both CR core promoters contain consensus TATA boxes at the preinitiation complex (PIC) location, while the TFIID core promoters contain a GA-element (GAE) at the PIC location. **B)** UAS-core promoter plasmid libraries are transformed into yeast with degron-tagged coactivator subunits (Taf13, TFIID; Spt7, SAGA; Med15, Mediator Tail). Nascent RNA barcodes are sequenced +/- auxin treatment, and UAS-core-barcode combinations are determined through sequencing of isolated plasmid libraries. Assay expression and log2-fold change after coactivator depletion are determined by normalizing RNA barcode counts to isolated plasmid counts. **C)** Violin plot comparing log2-assay expression of the *KRS1* (CR/TI) core promoter paired with all UAS sequences, compared to control regions from accessible chromatin (control acc.) or inaccessible chromatin (control inacc.). **D)** Violin plot of log2-assay expression of the *NOP13* (TFIID) core promoter, as in C.

### UASs broadly activate most yeast core promoters

We first compared the expression of UASs and negative control sequences fused to the four core promoters. For each of the four promoters tested, the median UAS activity is significantly higher than genomic control sequences regardless of whether these control regions are normally found in accessible or inaccessible chromatin (**Fig 1C-D, S1C-D**) (**Table S1**), indicating that these UAS regions activate transcription. For each core promoter, between 62%-74% of UAS sequences drive expression at a level higher than the median expression of negative control sequences. UASs showing low expression (**Fig 2A-B**) are above the non-UAS control expression median but below the control 90^th^ percentile, and High expression are UASs showing expression above the top 10% of non-UAS controls. 91% of all UASs drive expression higher than the median background in at least one core promoter context. Assuming a model of regulation in which gene expression is controlled in large part by the intrinsic activity of the UAS, we expect many UASs to display similar levels of activity across core promoters. Consistent with this model, we observe 40.1% of UASs driving expression above the control median at all four tested promoters (**Fig 2C**; termed generally active UASs), and a small subset of these generally active UASs drive high expression in all four core promoter contexts (9.2%, termed highly active UASs). These latter UASs are biased in being derived from CR genes (19.8% CR, compared to the 13.2% genomic average of CR genes, **Fig S2**). Relatively few UASs express below the genomic control median in all four promoter contexts (8.9%, termed non-active UASs). About half of all UASs do not fall into these defined classes but instead activate the four core promoters to varying degrees, and examination of the most highly promoter-specific UASs is addressed below. To test whether UAS function in the reporter assay reflects endogenous activity, we compared transcription of the reporter to transcription of the genes used to create the UAS library. Encouragingly, the median genomic expression level for the highly expressed and generally expressed UAS class genes are significantly higher than non-active UASs (**Fig 2D**). These results strengthen the connection between the reporter assay and the activity of the UAS sequences in their normal genomic contexts.

**Fig 2.**
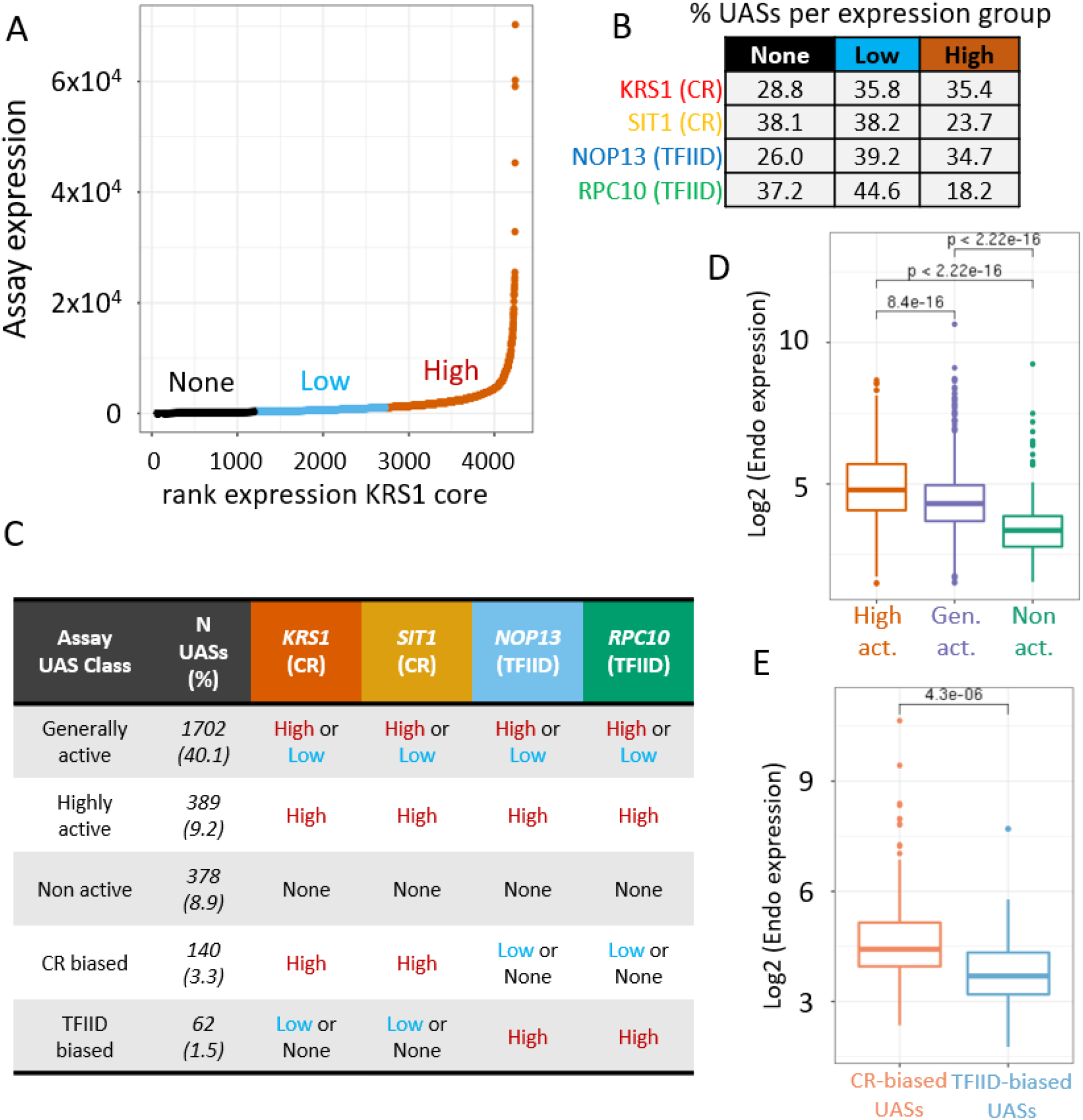
Expression categories of UASs in the UAS-core assay. **A)** Assay expression level vs. rank expression for all UASs paired with the *KRS1* core promoter, grouped by expression level relative to genomic controls. “None” expression group is expression below the median for genomic controls. “Low” expression group is higher than the median but below 90^th^ percentile for genomic controls. “High” is above 90^th^ percentile for genomic controls. **B)** Percentage of UASs in each expression group for UAS-core transcriptional assay per core promoter. Groups are defined as described in Fig2A. **C)** Classification of UASs based on expression groups as in Fig2A-B. Generally active UASs (40.1%) are in the “high” or “low” expression group for all four core promoters. Highly active UASs (9.2%) are a subset of the generally active UASs, and are in the “high” expression group for all four core promoters. Non active UASs (8.9%) are in the “none” expression group for all four core promoters. CR-biased (3.3%) and TFIID-biased (1.5%) UASs are in the “high” expression group only when paired with core promoters from their same regulatory class. **D)** Genomic expression level of genes with UASs defined in the UAS-core assay as highly active (High act.), generally active (Gen. act.), or non active (Non act.). **E)** Genomic expression level of genes with UASs defined in the UAS-core assay as CR-biased of TFIID-biased.

To determine whether specific regulatory factors are associated with our defined UAS classes, we performed sequence motif enrichment analysis using Homer^31^ (**Table S2**). In the highly active UAS class, we observed a significant enrichment of DNA binding motifs for transcription factors Sfp1, Rap1, and Reb1. Sfp1 is known to regulate ribosomal protein genes (RP) and ribosomal biogenesis genes (RiBi)^32,33^, and Rap1 associates with many genomic loci including highly expressed ribosomal genes^34,35^. RiBi and RP genes are overrepresented in the highly expressed class of UASs (3.5-fold and 5.1-fold above the genome average, respectively). Reb1 is a general regulatory factor known to drive strong expression at a variety of protein coding genes^36,37^, and its motif is also enriched in the generally active UAS class. Also modestly enriched are motifs for the general regulatory factor Abf1 and the RP gene regulators Fhl1 and Tod6 (**Table S2**)^37–39^. Our results suggest that these general regulatory factors can broadly activate core promoters regardless of promoter class.

### Differences in regulation between TFIID and CR promoters

Although the above results suggest that a substantial proportion of UASs have intrinsic activity, about half of the UASs don’t appear to generally activate all four core promoters. We next focused on the behavior of the most highly specific UASs. For the small subset of UASs that display highly biased expression at CR core promoters (n = 140, **Fig 2C**), the median genomic expression of the endogenous genes controlled by those UASs is significantly higher than the genomic median for TFIID-biased UASs (N = 62 **Fig 2E**). Furthermore, CR-biased and TFIID-biased UASs are more frequently derived from CR and TFIID-class genes, respectively (**Fig S2A**), indicating that this subset of UASs display class-specific matching with compatible endogenous core promoters. No sequence motifs were found enriched in either class of highly specific UAS sequences, and genes containing these UASs on average display the same level of responsiveness to MED Tail, SAGA and TFIID depletion compared to CR or TFIID UASs not found to display high degrees of core promoter specificity (**Fig S3**).

Prior work demonstrated that inducible metazoan core promoters tend to exhibit lower intrinsic activity than constitutive core promoters^40,41^. To test if this feature is conserved in yeast, we examined whether CR promoters display low intrinsic activity, given that CR genes, like metazoan inducible genes, tend to be stress responsive and are enriched for TATA-containing promoters. Examining the range of expression levels promoted by the complete set of UASs, we observed that the two CR promoters (both the MED Tail-dependent and independent promoters) on average display a lower intrinsic activity and, as a class, display a higher dynamic range of expression than the two TFIID promoters (**Fig S1E**). For example, the lowest expressed UAS-CR promoter combinations are well below the lowest expressed UAS-TFIID promoter constructs, and these expression differences between TFIID and CR promoters disappear in the most highly expressed UAS-promoter constructs (**Fig S1E**). Further supporting the general trend of low basal activity and high inducibility, CR core promoters display a higher fold-induction when paired with their genomic UASs (**Fig S1F**). Together, our findings suggest that lower intrinsic activity of many inducible core promoters is a conserved feature in eukaryotes.

Although few UASs display strict specificity for core promoters, it’s possible that the quantitative activity of UAS elements depends on matching with a core promoter of the same gene class. For example, individual UASs derived from many CR genes can activate transcription from both CR and TFIID core promoters but may promote the highest expression when paired with a CR promoter. Previous findings in mammalian cells^23^ suggested that expression of most enhancer and promoter combinations can be explained by the combined intrinsic activities of enhancer and promoter, and that enhancer-promoter combinations expressing higher than expected, based on their intrinsic activities, tend to be matched by regulatory class.

To determine whether deviations from intrinsic UAS activity correspond to coactivator class-specific regulation, we examined the correlation in expression for all UASs driving expression from the same core promoter class (*KRS1* vs. *SIT1*) or from core promoters of different class (*KRS1* vs. *NOP13*). We found that the correlation for UASs at the same core promoter class is slightly higher (R^2^(adj) = 0.42, CR core vs. CR core, **Fig3A**) than for UASs in differing promoter class context (R^2^(adj) = 0.31, CR core vs. TFIID core **Fig3B**), suggesting subtle expression differences driving different class promoters. We next asked if these observed differences in UAS activity at different core promoters correlate with the gene class the UAS was derived from. Strikingly, we observe that UASs that more strongly activate CR promoters compared to TFIID promoters (**Fig 3C**, blue bars, linear regression residual quartiles 3-4) are more frequently derived from CR genes. We observe a slight trend where UASs more strongly activating TFIID promoters (**Fig 3C**, blue bars, linear regression residual quartiles 1-2) are more frequently derived from TFIID genes. These trends are not apparent when comparing core promoters of the same class (**Fig 3C**, greyscale bars, CR vs. CR or TFIID vs. TFIID). In addition, UASs displaying lower activating activity at CR class promoters display higher variance across the four core promoters (**Fig S2B-C**), a trend not apparent at TFIID promoters, suggesting this specificity is in part due to functional incompatibility of UASs with core promoters for the CR class. Our results are consistent with a model in which the intrinsic activity of UASs on all four promoters generally correlates with expression level of the endogenous gene, but deviations from this model can be explained in part through matching of regulatory elements from the same coactivator class.

**Figure 3:**
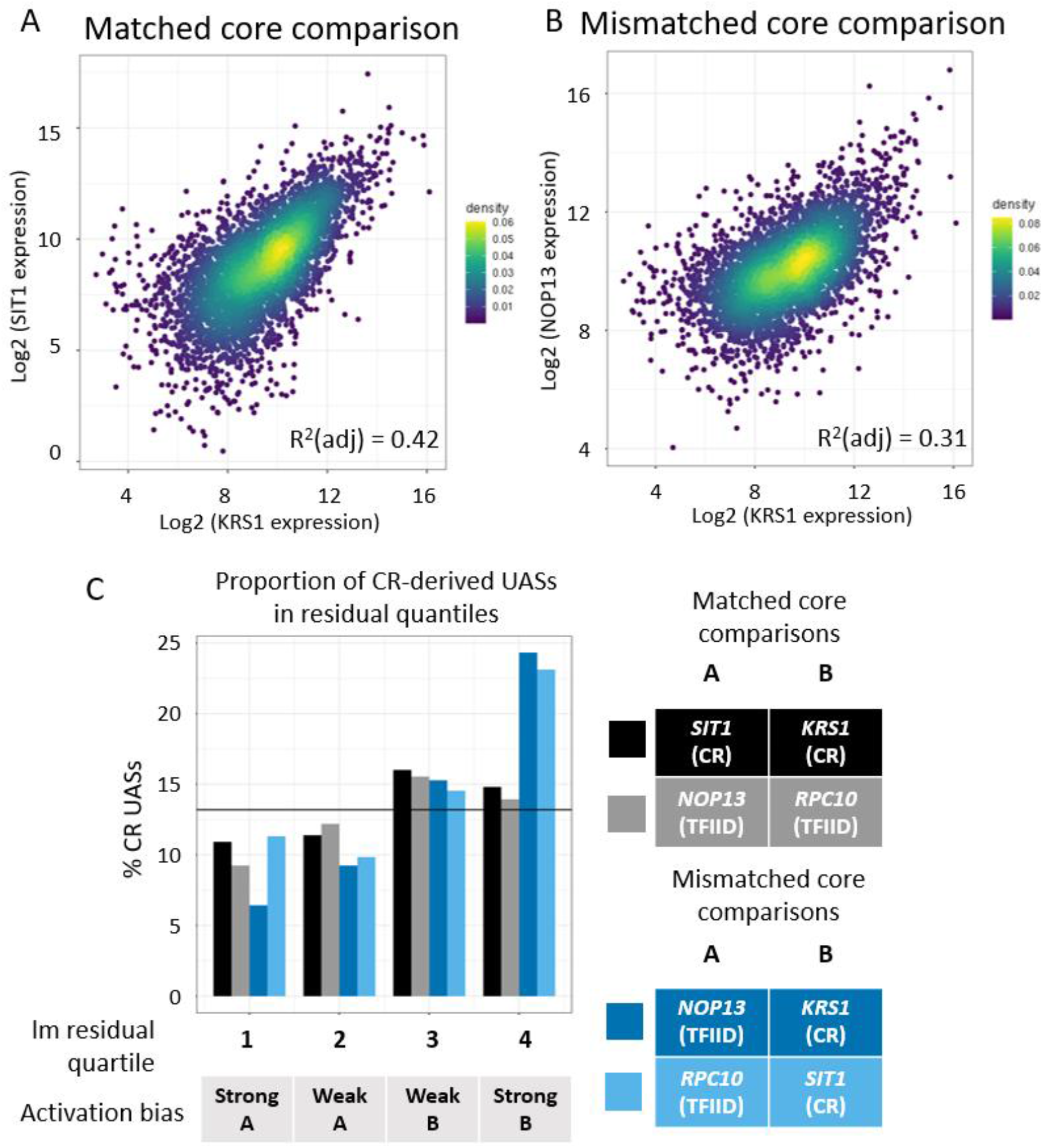
Differences in expression level for UASs in different core promoter contexts correlates with coactivator class. **A)** Scatter plot comparing log2-expression of UASs in *KRS1* (CR) promoter vs. *SIT1* (CR) promoter. Correlation displayed is the adjusted R-squared from a linear regression model (lm). **B)** Scatter plot comparing log2-expression of UASs in *KRS1* (CR) promoter vs. *NOP13* (TFIID) promoter. **C)** Linear regression model (lm) residuals were calculated for each UAS in matched core comparisons (CR-CR and TFIID-TFIID) or mismatched core comparisons (CR-TFIID). The proportion of UASs from CR genes is plotted for each residual quantile for the four comparisons shown. Black line represents genomic CR proportion (13.2%).

### SAGA and MED Tail specificity are dependent on both UAS and core promoter, while TFIID is primarily dependent on core promoter

We next asked whether dependence on particular cofactors is specific to the core promoter sequence, UAS sequence, or a combination of both. For these studies, we examined the change in transcription for UAS-core combinations following rapid depletion of Taf13 (TFIID), Spt7 (SAGA) or Med15 (MED Tail) and assessed whether there are differences in coactivator sensitivity dependent on UAS class. We found that CR promoters on average are significantly more Med15 sensitive when paired with UASs from CR genes (**Fig 4A**), while TFIID core promoters on average display no sensitivity to Med15 depletion regardless of the UAS. These findings suggest that functional matching of UAS and core promoter sequences is important for dependence on MED Tail. Similarly, CR core promoters on average are more sensitive to Spt7 depletion when paired with CR UASs (**Fig4B**). In contrast, TFIID promoters are generally more sensitive to Taf13 depletion vs CR promoters (**Figs 4C, S4C**) and no difference in Taf13 dependence was observed for TFIID promoters paired with CR vs TFIID UASs. This suggests that TFIID sensitivity is primarily dependent on core promoter sequence. Taken together, our results indicate that the UAS and core promoter both play significant roles in SAGA and MED Tail dependence, while the core promoter is the major driver of TFIID dependence.

**Figure 4:**
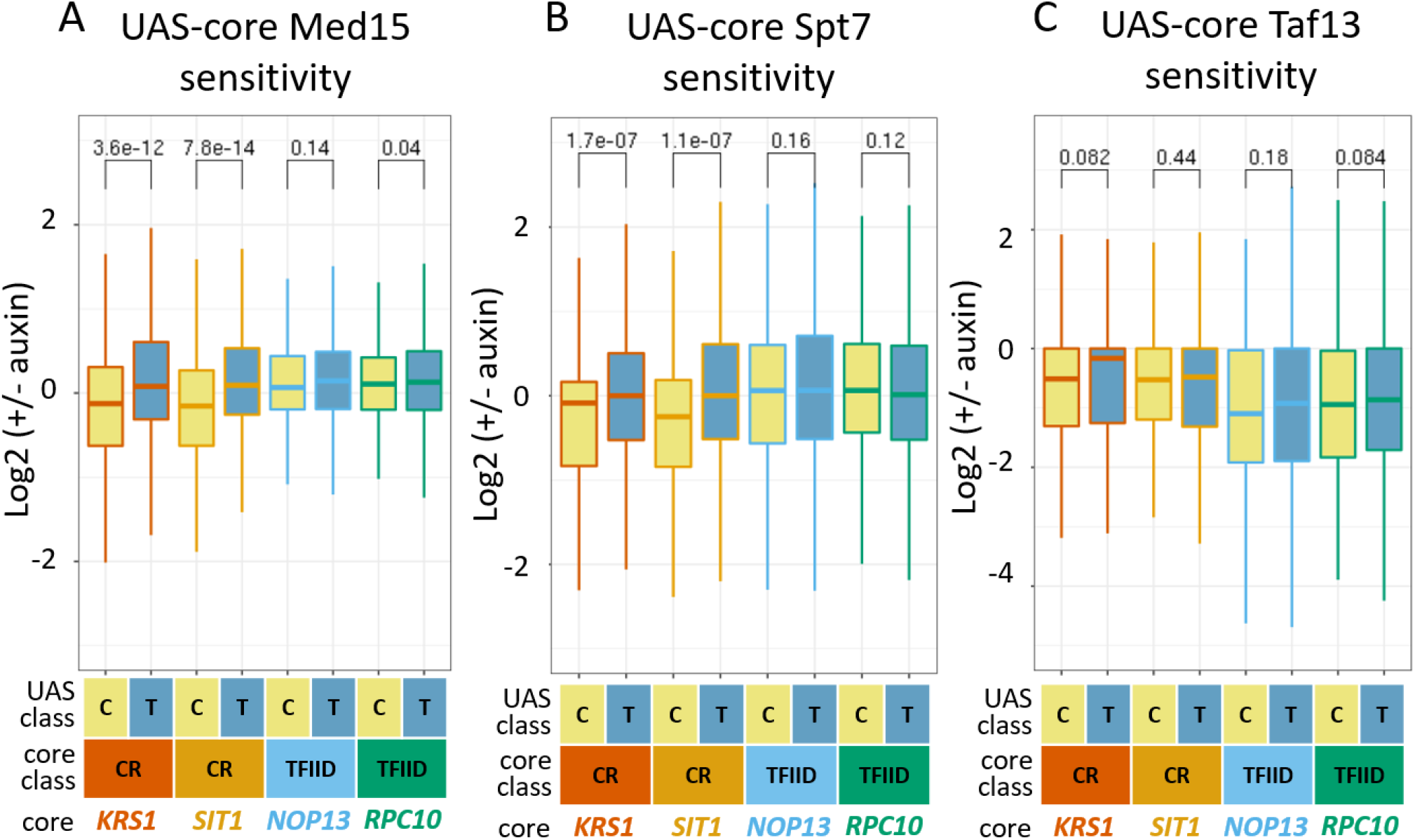
Coactivator sensitivity dependence on UAS and core promoter classes. **A)** Log2-fold change in transcription after Med15 depletion for the four test core promoters, comparing UASs from CR class genes (C) or TFIID class genes (T). T-test statistic is shown. **B)** Comparison as in A, but assessing log2-fold change in transcription after Spt7 depletion. **C)** Comparison as in A-B, but assessing log2-fold change in transcription after Taf13 depletion.

### Roles of the UAS and TATA in Mediator Tail dependence

Given that only half of CR genes are Tail-dependent, we examined sensitivity of the UAS-promoter constructs to rapid Med15 depletion. Interestingly, when paired with UASs from MED Tail-dependent genes, the core promoter from *SIT1* (a Tail-dependent gene) appears more sensitive to Med15 depletion compared to the core promoter of *KRS1* (a Tail-independent gene) (**Fig 5A**). Since both promoters contain TATA, it suggests that additional features in the core promoter may contribute to MED Tail sensitivity.

**Figure 5:**
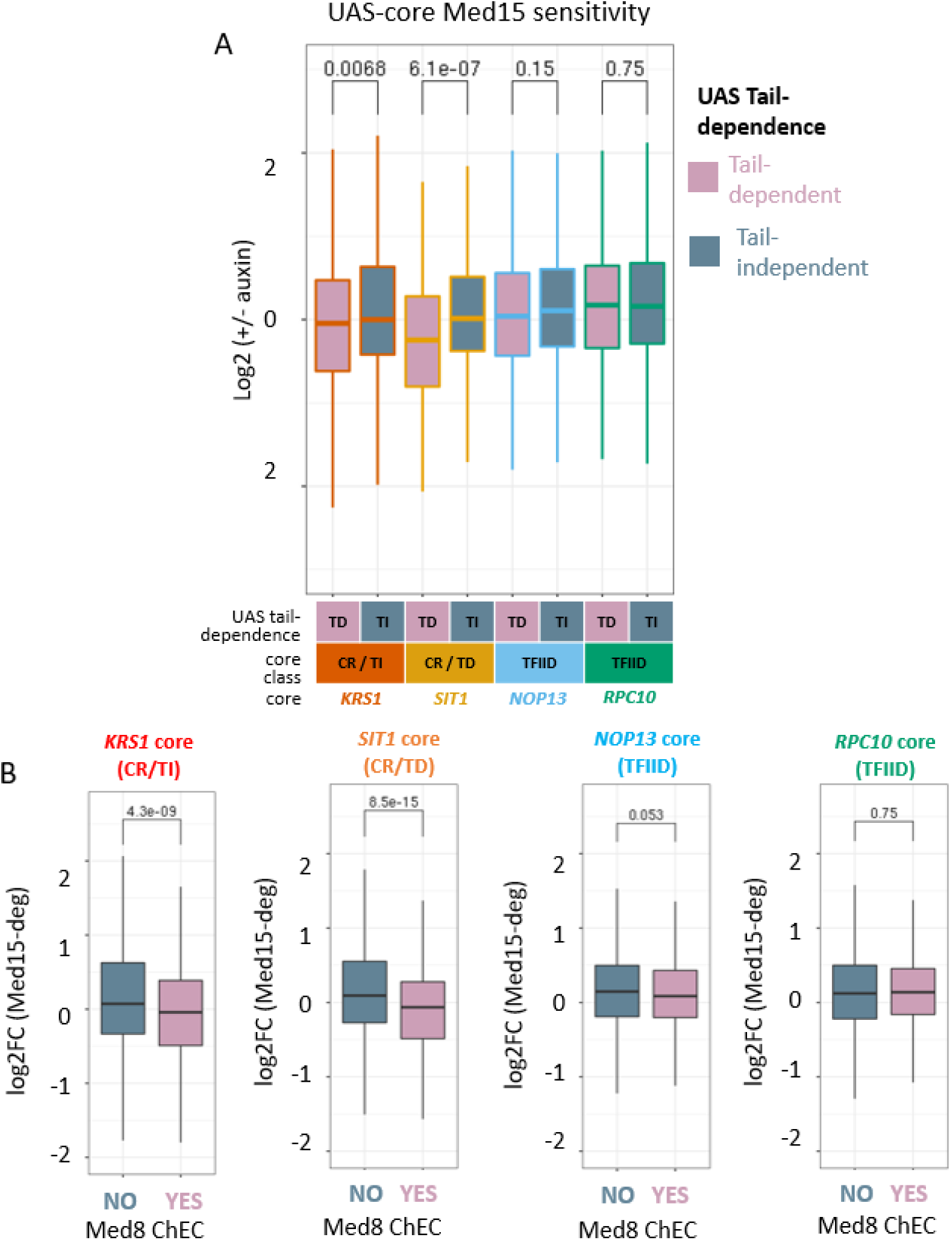
Mediator Tail sensitivity is highly dependent on UAS localization and core promoter identity. **A)** Log2-fold change in transcription after Med15 depletion for the four test promoters, comparing UASs from Mediator Tail-dependent vs. Tail-independent genes. Tail dependence of CR cores is noted (TI = Tail-independent, TD = Tail-dependent). T-test statistic is shown. **B)** Log2-fold change in transcription after Med15 depletion for the four test promoters, comparing genes with endogenous Mediator ChEC-seq signal in the UAS region vs. genes without endogenous Mediator ChEC-seq signal.

Recently, we mapped MED binding using the MNase-based ChEC-seq assay and found that MED is detected at 16.6% of genomic UASs (~70% of CR UASs and ~18% of TFIID UASs) while expression from only ~6.5% of genes, nearly all from the CR gene class, are MED Tail-dependent^18^. Our analysis here shows that, on average, CR core promoters are more sensitive to Med15 depletion when paired with a UAS having detectable MED binding in the genome, whether this UAS was derived from a CR or TFIID gene (**Fig 5B**). These results imply that Tail dependence is linked to the ability of factors bound at the UAS to recruit MED. In contrast, TFIID core promoters do not show Med15 sensitivity regardless of detectable MED as the UAS, confirming the above suggestion that the core promoter is also an important component of MED Tail sensitivity.

Our results above indicate that the core promoter plays an important role in MED Tail sensitivity so we further analyzed core promoters of Tail dependent genes. 60% of CR promoters contain a TATA motif (TATAWAW), and MED Tail-dependent promoters are further enriched for the TATA motif (70%). One hypothesis is that, for the minority of TATA-less Tail-dependent genes, additional core promoter elements contribute to Tail sensitivity. We performed sequence enrichment analysis of core promoters lacking annotated TATA boxes that were derived from either Tail-sensitive or Tail-insensitive genes. Surprisingly, the top motif enriched in the Tail-sensitive promoters is a TATA-like element (TWTWWA), which is found within 65 bp upstream of the TFIIB binding site in 74% of Tail-sensitive TATA-less promoters. CR genes with canonical TATA motifs in their core promoters are moderately sensitive to Med15 depletion, while CR genes with TATA-like elements are weakly sensitive to Med15 depletion (**Fig 6A**). Endogenous CR genes without any TATA or TATA-like element are on average insensitive to Med15 depletion. Endogenous SAGA sensitivity appears high for all CR class genes regardless of core promoter element but appears highest with a canonical TATA box (**Fig 6B**). Similar weak TATA boxes were previously found to moderately increase transcription in yeast synthetic promoters^42^, an effect likely due to moderate engagement with the MED Tail pathway based on the results described here.

**Figure 6:**
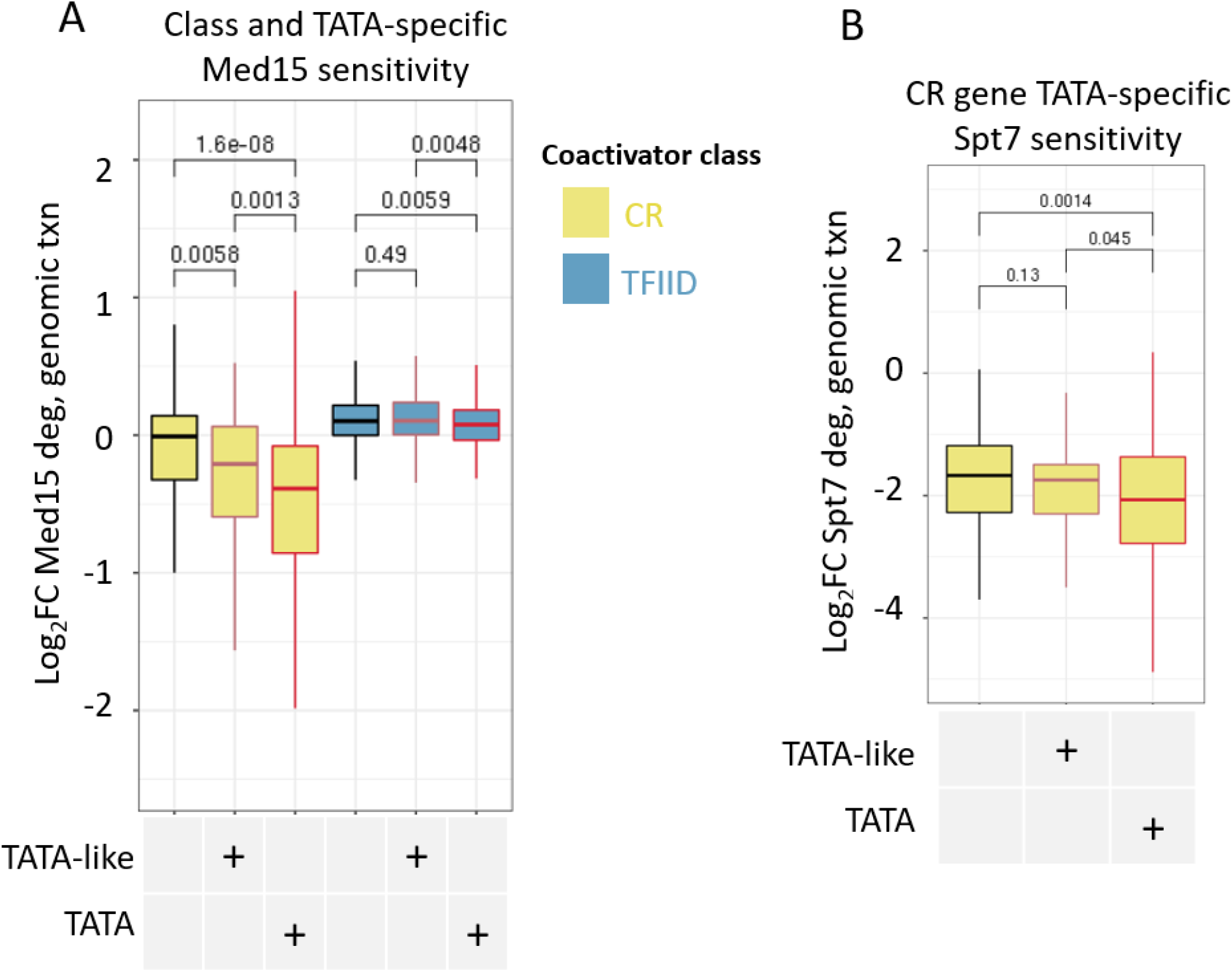
TATA-like elements and Mediator UAS localization independently promote Mediator Tail sensitivity. **A)** Log2-fold change in genomic transcription after Med15 depletion (Data from Warfield 2022), comparing CR class vs. TFIID class genes, additionally comparing genes containing a TATA-like element (TWTWWA), canonical TATA box (TATAWAW), or no TATA element in the core promoter. T-test statistic is shown. **B)** Log2-fold change in genomic transcription after Spt7 depletion (Data from Donczew 2020), comparing CR class genes containing a TATA-like element (TWTWWA), canonical TATA box (TATAWAW), or no TATA element in the core promoter.

**Figure 7:**
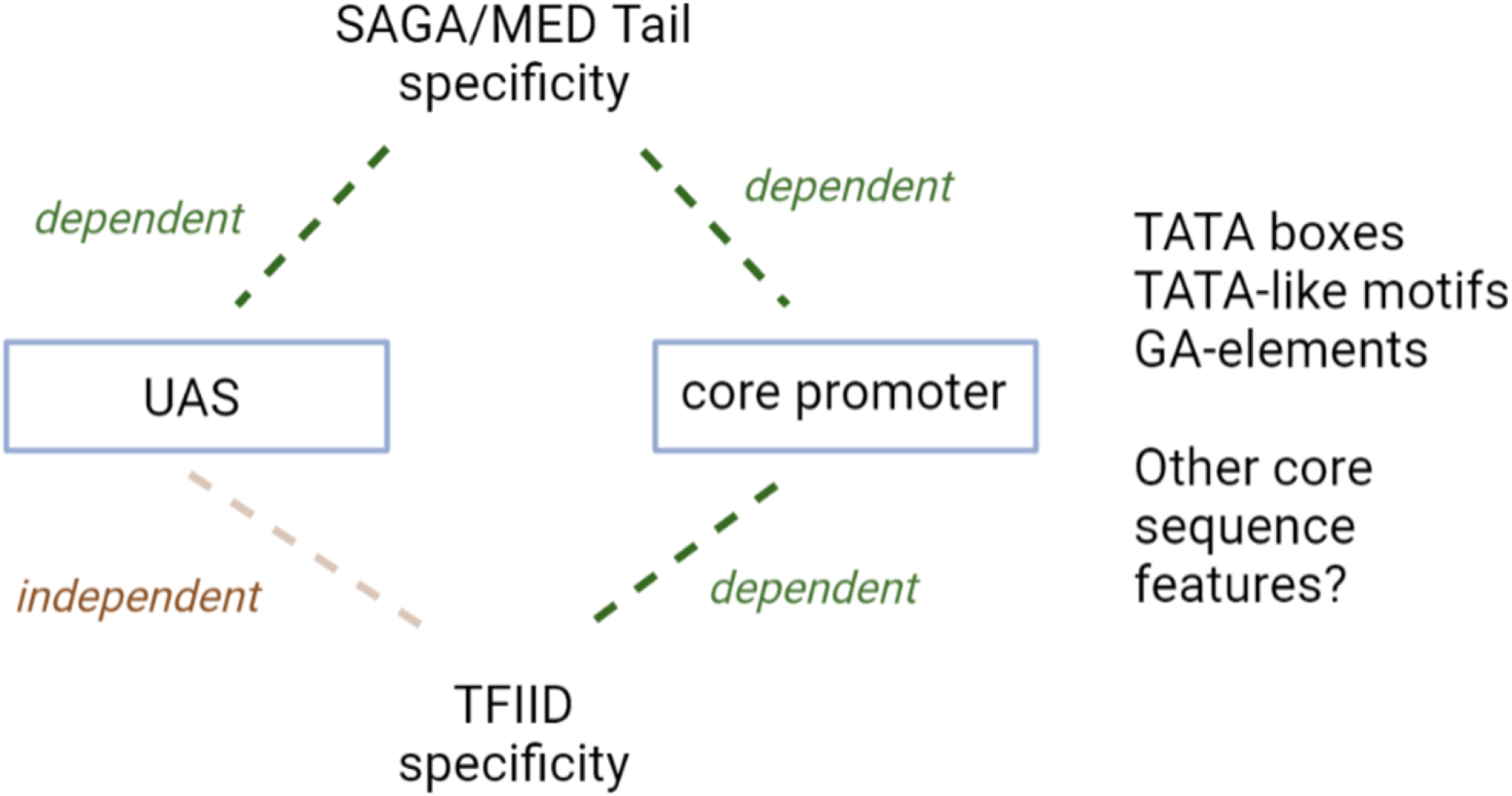
model. Mediator Tail and SAGA sensitivity depend on a combination of UAS localization and flexible core promoter elements. TFIID sensitivity is dependent on the core promoter sequence.

## Discussion

Here we describe a large-scale reporter assay to assess the functional matching of yeast UAS and core promoter sequences as well as the basis for coactivator specificity that defines different gene classes. First, we found that a large proportion of UASs can activate core promoters irrespective of gene class. This suggests that most UASs do not show strict specificity for activation of yeast core promoters and, importantly, that most yeast UASs display intrinsic activity. The low absolute specificity of UASs is in contrast to properties observed in *Drosophila*^22^, in which enhancers from housekeeping and developmental gene classes have been observed to display high selectivity for promoters from the same regulatory class. This dichotomy may be due to *Drosophila* transcription factors that are known to preferentially bind at the two classes of regulatory enhancers and to several common consensus motifs that are found at many *Drosophila* core promoters. We did find a low percentage of UASs (<5% total) that are highly specific for TFIID or CR promoters, but these are the exceptions to the general rules of broad compatibility, and no specific regulatory factors appear enriched in these class-specific UASs, based on the sequence motifs of known transcription factors.

While only a small number of UASs show strict specificity, we found that matching of UASs and core promoters by class is important in dictating expression output. For example, transcription of the UAS-core constructs is significantly higher with matched UASs and core promoters from the same gene class. These latter results are consistent with recent findings in both human^23^ and mouse cells^24^, in which the transcriptional output of enhancer-promoter combinations generally correlates with the intrinsic activities of enhancers and core promoters, although class-matched regulatory elements display higher expression levels. These combined results suggested that functional compatibility contributes to the interplay between mammalian enhancers and promoters.

The basis for specificity of SAGA and MED Tail function is very likely mediated by the sequence specific transcription factors targeted to the UAS. For example, our results show that UASs which recruit detectable MED in their normal chromosomal locations are preferentially Tail-dependent when paired with an appropriate core promoter. ~18% of TFIID dependent genes are linked to UASs that bind MED. However, the incompatibility of the core promoter with Tail-sensitivity explains why these genes are normally insensitive to rapid Tail depletion. It is likely that the MED-recruiting UASs are bound by activators that directly interact with MED Tail, the only known activator binding domain in yeast MED. In contrast to the recruitment of MED that is limited to a subset of UASs, SAGA is recruited to a high percentage of UASs in both CR and TFIID class genes^16,43^, yet SAGA dependence is confined to CR genes. Thus, SAGA-specificity is also in part dictated by the core promoter – a result confirmed by our reporter assays.

Our findings show that TFIID sensitivity is largely specified by the core promoter sequence, consistent with metazoan TFIID subunits that are known to interact with core promoter elements^44^. However, our work suggests that the UAS plays an important role in the transcription of these TFIID-class genes. Recent work mapping transcription factors in yeast concluded that many TFIID-class genes are devoid of transcription factor binding with the exception of insulator factors (mainly Reb1 and Abf1) and are constitutively transcribed^45^, suggesting that many TFIID-class core promoters would be unresponsive to TFs. In contrast, our assays show that TFIID-class core promoters are responsive to a broad range of UASs, albeit with a lower dynamic range than the TATA-containing CR promoters. Interestingly, we found that the small fraction of highly TFIID-biased UASs is normally associated with TATA-less and TFIID-class genes, suggesting that functional compatibility between UAS and promoter is a feature of these highly specific elements.

In this study we found that both UAS and core promoter are vital contributors to coactivator specificity. Although individual transcription factors are known to interact with MED-Tail, it remains unclear which specific factors or combinations of factors binding to the UAS are responsible for coactivator specificity. Emerging evidence suggests that TF binding does not always correlate with transcription^46,47^, therefore functional mapping of transcription factors across the yeast genome will be necessary to determine the roles of specific TFs in promoting coactivator specificity at any particular gene. In addition, core promoter motifs other than the TATA box likely play important roles in coactivator specificity, and often large-scale experimentation and machine learning is necessary to identify these elements^48^. Testing additional core promoter sequences in this or similar assays will reveal expanded roles of the core promoter in gene-specific transcriptional regulation.

## Limitations of the study

Our assays were performed using a reporter plasmid library and it is known that the genomic environment affects the absolute level of transcription. Based on prior results, it is likely that the absolute but not relative expression level of UAS-core combinations is partially dependent on chromatin^40,49^, so use of the reporter likely does not affect our conclusions on functional compatibility and coactivator specificity.

The core promoters selected correspond to a small selection of genes displaying moderate endogenous expression to be consistent in reporter expression. There are likely differences in how promoters with high or low intrinsic activity respond to UASs and coactivators, therefore testing additional core promoters will expand our understanding of the functional connection between regulatory elements.

## Methods

### Resource availability

#### Lead contact

Further information and requests for resources and reagents should be directed to and will be fulfilled by the lead contact, Steven Hahn.

#### Materials availability

All unique/stable reagents generated in this study are available from the Lead Contact without restriction.

### Data and code availability

- The datasets generated during this study are available at Gene Expression Omnibus under accession XXXXXX, Reviewer Token YYYY.
- All code snippets and whole notebooks are available from the Lead Contact upon request.
- This paper does not report original code.
- Any additional information required to reanalyze the data reported in this paper is available from the lead contact upon request.

### Experimental Model and Subject Details

All *S. cerevisiae* strains used in these studies are derived from strain BY4705 (Brachmann et al., 1998) with genotypes listed in **Table S3**. Genotypes and source of *S. pombe strains* used as spike-in controls for RNA-seq are also listed in **Table S3**. Culture conditions are listed in Method Details.

#### Defining regulatory regions

Core promoter regions for yeast genes were defined by the location of the preinitiation complex, inferred by the location of TFIIB ChIP-seq signal. TFIIB ChIP-seq from several sources^45,50^ was used to define a consensus TFIIB location. Consensus TFIIB locations located downstream of the annotated transcription start site, overlapping with the annotated coding region, or further than 150nt upstream of the annotated transcription start site were removed from our list of core promoters, resulting in 4273 genes for inclusion in the assay. The upstream edge of the core promoter was defined as 60nt upstream of the TFIIB location, which encompasses most TATA elements for both CR-class and TFIID-class genes (a minority of TFIID-class genes contain TATA boxes) but minimizes regions of upstream activator binding. UAS regions were defined as 200nt directly upstream of the TFIIB-defined core promoter, and were cross-referenced with available ChIP-exo data^45^ to ensure these UAS regions are able to bind TFs. A majority of activator and general transcription factor binding sites near genes included in this assay overlap with these 200nt UAS regions (i.e. 86% of Hap1, 81% of Lys14, 84% of Met31, 86% of Yap1, 87% of Abf1, and 88% of Reb1 binding sites). Theoretical extension of these regions by an additional 50nt increases binding sites by these factors by less than 5% on average, validating that a vast majority of factor binding is contained within the 200nt region.

Random sequence controls sequences 200nt in length were sampled from the genome to include an even number of sequences from accessible chromatin (defined by FAIRE-seq signal^51^) and inaccessible chromatin, as well as sequences in coding regions and non-coding regions. For each selected sequence, its reverse complement was also included, for a total of 224 control sequences.

Four representative core promoters were selected for the UAS-core assay to include ranging regulatory behavior: two core promoters from CR class genes containing TATA elements, one Mediator Tail-dependent (*SIT1*) and one Tail-independent (*KRS1*); and two core promoters from TFIID class genes containing GAE elements (*NOP13* and *RPC10).* All genes display moderate expression levels in yeast and, to confirm expression in the reporter context, all four core promoters were cloned into the reporter with and without their respective UASs and expression quantitated by RT-qPCR.

#### Cloning of combinatorial UAS-promoter plasmid libraries

Reporter plasmids were derived from the Yeast_Dual_Reporter plasmid (<u>Adgene, Plasmid 127546</u>) in which the GFP region was replaced with an unstable version of GFP^52^. Plasmid libraries were cloned using Gibson assembly of three fragments: a UAS sequence library (~16,000 sequences, Twist Biosciences) containing 200nt UAS sequences from the above analysis and 25nt of homology to the reporter plasmid (upstream of UAS) and 25nt of homology to one of the four core promoters (downstream of UAS); a core promoter fragment (Integrated DNA Technologies) containing a G-less randomized 18nt barcode in the 5’UTR of uGFP with homology to the reporter plasmid (downstream of core); and a ~10kb fragment containing the remainder of the reporter plasmid with bacteria and yeast selection markers (**Table S4**). Plasmid libraries were generated for each core promoter individually: Four Gibson reactions were pooled and ethanol precipitated, and this pool was transformed into electrocompetent DH10b *E. coli* (8 transformations per pool). These transformations were grown overnight in 50ml YT+Amp, and plasmid libraries were purified using the Qiagen Midi-prep kit. The complexity of each core promoter plasmid library was estimated by plating a portion of the overnight culture, counting the number of colonies from this portion and extrapolating to the total culture, and ranges from 16-44 randomized barcodes per UAS-core sequence combination. The four core promoter libraries were pooled according to their estimated complexities.

#### Yeast strains, transformation, and culturing

Med15, Taf13, and Spt7 degron strains are derivatives of BY4705^53,54^, and construction of these strains is described in Donczew et al. 2020^16^ and Warfield et al. 2022^18^ (**Table S3**). The pooled plasmid library was transformed into each degron strain using the lithium acetate method. The efficiency of transformation was estimated by plating a portion of the culture and was found to be at least tenfold higher than the plasmid library complexity for each sample. Transformants were grown at 30C in 200 ml -Ura synthetic complete media (SC) and were split 1:4 in fresh -Ura SC each time the culture reached an OD 600nm above ~1.5. After 3 splits, the culture was diluted to an OD of 0.2 for final growth. At mid-log phase (OD ~ 0.8), a 25 ml sample of each culture was treated with 500 μM 3-IAA (auxin) in DMSO to rapidly degrade coactivator subunits, and a control sample was treated with DMSO alone. After 30 min of +/- auxin treatment, all samples were treated with 4-thiouracil (5 mM) for 5 min to label nascent RNAs. All samples were pelleted by centrifugation, flash frozen and stored at −80 C until further processing. A total of 4 replicates of +/- auxin were performed for each coactivator degron strain in 2 separate batches beginning from library transformation.

#### Nascent RNA isolation

Total RNA was extracted from stored pellets using the Ribopure kit with the addition of 10mM beta-mercaptoethanol to buffers RLT and RPE, and using care to keep 4-TU containing RNA protected from light. Nascent RNAs were enriched according to previous protocols^55^: briefly, 40 μg of input RNA was biotinylated using MTS-biotin. The biotinylation reaction was purified using phenol/chloroform/isoamyl alcohol, followed by ethanol precipitation of RNA from the aqueous phase. Nascent RNAs were captured using streptavidin beads (Dynabeads MyOne Streptavidin C1) with high salt washes to reduce non-specific RNA binding to beads. Nascent RNAs were eluted from the beads using a reducing elution buffer, and cleaned up using an RNeasy Minelute kit. Nascent RNA was quantified using the Qubit High Sensitivity RNA assay.

#### Plasmid library isolation

Plasmid libraries were isolated from -Ura selected yeast sampled immediately prior to auxin treatment. A modified Qiagen Mini-prep protocol (see Qiagen online resources) was performed on yeast pellets from 15 ml of culture. Briefly, yeast pellets were resuspended in buffer P1, transferred to tubes containing 100 μl beads, and disrupted using a bead beater for 5 min. Tubes were spun briefly, and the lysate was transferred to a fresh 1.5 ml Eppendorf tube. The remainder of the standard Mini-prep protocol was followed, typically yielding 5-10 ng/μl of plasmid libraries.

#### RNA barcode library preparation

cDNA was generated from purified nascent RNAs using the <u>Transcriptor Reverse Transcription kit</u> and a custom RT primer complementary to the uGFP reporter gene. After reverse transcription, cDNA was purified using ethanol precipitation, and was amplified with 4 rounds of PCR using targeted primers containing anchor sequences for indexing. Amplified cDNA was purified, and size selected using <u>Biomag SPRI</u> beads, and qPCR was used to determine the number of amplification steps necessary for index PCR. Dual-indexed libraries were generated with a second PCR reaction using unique indexing primers complementary to PCR 1 anchors, purified and size selected using <u>Biomag SPRI</u> beads, and pooled according to Tapestation quantification.

#### UAS-core-barcode library preparation

To identify the UAS and core promoter sequences associated with observed RNA barcodes, sequencing libraries were prepared from isolated reporter plasmid libraries. 10 ng of plasmid libraries were amplified with 4 rounds of PCR followed by size selection and qPCR quantification as outlined above. Dual-indexed libraries were generated with a second PCR reaction using unique indexing primers complementary to PCR 1 anchors, purified and size selected using <u>Biomag SPRI</u> beads, and pooled according to Tapestation quantification.

#### Sequencing

RNA and DNA libraries were pooled, and a final library concentration was determined through qPCR. Paired-end 100 nt sequencing was performed on the <u>NextSeq P2</u> platform. Due to the relatively low complexity of reporter plasmid regions in targeted sequencing, a 50% PhiX spike-in was performed.

#### Sequence processing

RNA barcode and DNA plasmid library read pairs were joined into amplicons. To account for mutations in sequencing, amplicons were clustered into representative sequences using VSEARCH^56^, generating clusters of 99% similarity for RNA barcode amplicons and 97% similarity for DNA plasmid amplicons. Targeted nascent RNA barcode cluster counts were highly reproducible between replicates (Pearson range 0.84-1.00). Cluster outputs were converted to tables containing consensus sequences and counts for plasmid UAS-core-barcode regions and expressed RNA barcodes (R, package Tidyverse^57^). The identities of UAS and core promoter regions were determined using a distance join function to a table of reference sequences (package fuzzyjoin^58^). RNA barcodes were matched to UAS-core combinations using an exact join. The following filtering was performed: within each sample replicate, the most common DNA cluster sequence for each barcode was identified, and all other DNA sequences per barcode were filtered out to eliminate background in sequencing. Next, across all samples, any barcodes observed in more than one UAS-core combination were removed to eliminate ambiguity in the source of barcode expression. After data processing and filtering, the average number of unique barcodes observed per UAS-core combination was 12. Per-sample DNA and RNA counts for each UAS-core combination was determined by summing counts for all unique barcodes assigned to that UAS-core combination. Final expression values for each UAS-core combination per sample were determined by normalizing RNA counts and DNA counts to each sample’s library size, then dividing normalized RNA counts to normalized DNA counts. These normalized barcode counts are also highly reproducible (Pearson range 0.71-0.97). Coactivator degron fold change values were determined by experimental batch, calculated as the log median expression for auxin treated samples divided by DMSO treated samples. R plots were generated using R packages ggplot2^59^ and viridis^60^. T-tests for boxplots were performed using ggpubr^61^.

#### Motif analysis

Differential motif enrichment was performed using Homer^31^ (findMotifs) by comparing UAS sequences in each defined class against a background of all remaining UAS sequences using default parameters.

#### Variance calculation

The variance for each UAS (see Fig S2B-C) was calculated as the sum of the absolute difference in UAS activation for each UAS-core combination and the mean UAS expression across the four core promoters, divided by the mean UAS expression across the four core promoters.

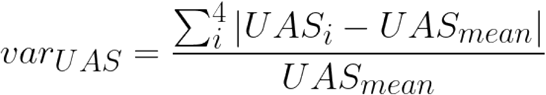

#### Quantification and Statistical Analysis

All next-generation sequencing experiments were done in four biological replicates. P-values for mean comparisons displayed in boxplots were determined using a T-test. The details of quantification are included in the following sections of the Methods: “Sequence processing”. The details of statistical analysis applied are included in the following sections of the Methods: “Motif analysis”, “Variance calculation”.

## Acknowledgements

J.A.S. was supported by NIH T32CA009657. J.A.S. and S.H. were supported by NIH 6R35GM140823. This work was also supported by P30 CA015704 Cancer Center Support grant to FredHutch genomics. We thank FredHutch Genomics and Bioinformatics core for sequencing and consultation, especially Elizabeth Jensen, Dolores Covarrubias, Andy Marty, and Matt Fitzgibbon. We thank members of the Hahn lab and Toshio Tsukiyama for helpful discussions on this manuscript. We thank E. Ünal for providing the plasmid containing the unstable GFP variant.

---

**Fig S1.**
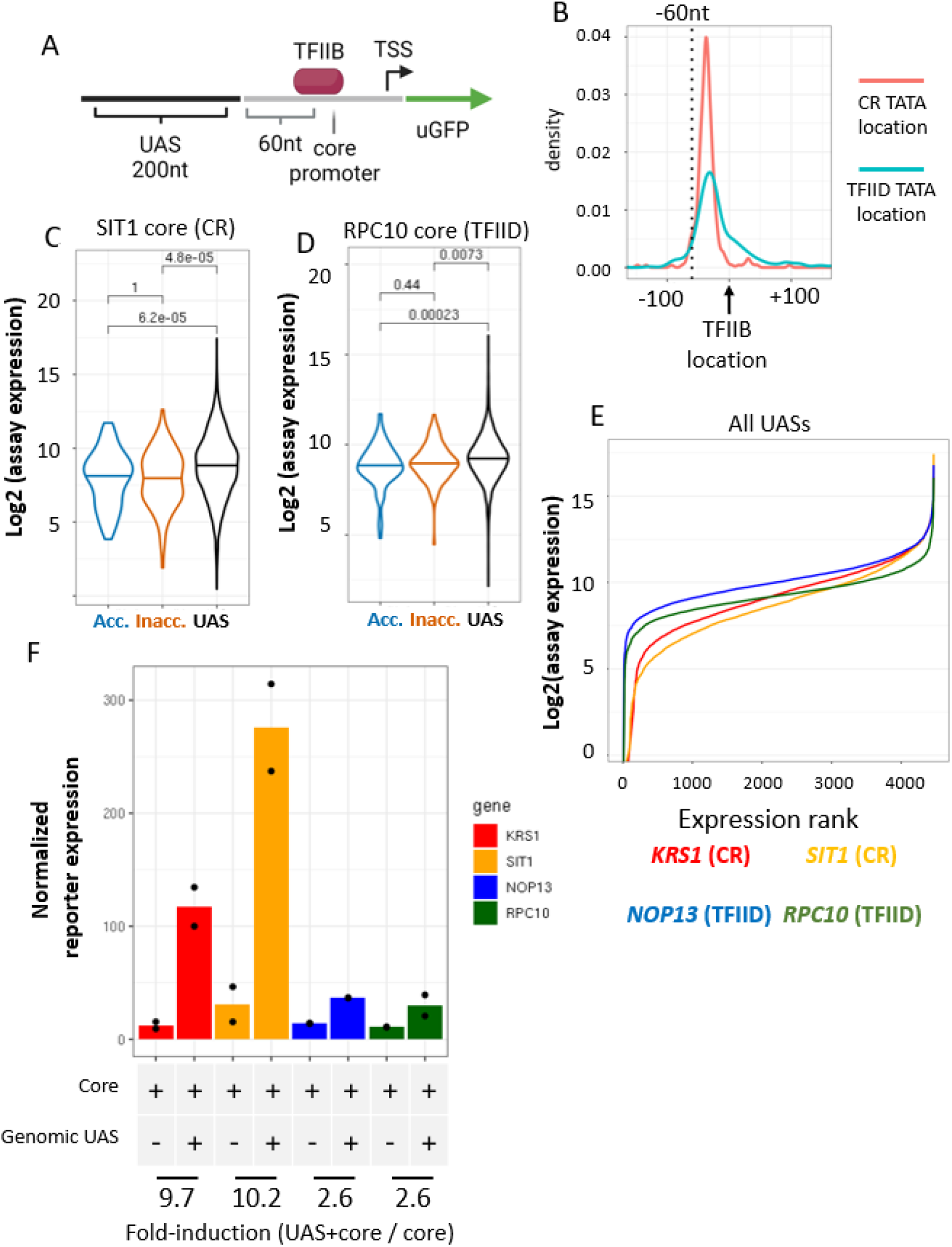
**A)** Scheme of UAS and core promoter definitions. Upstream edges of core promoters are 60nt upstream of the peak TFIIB ChIP-seq locations, and downstream edges contain the TSS. UASs are the 200nt region upstream of the defined core promoter region. **B)** Density plot of annotated TATA box locations relative to TFIIB locations for both CR and TFIID class genes. A majority of TATA boxes are located within the 60nt upstream of TFIIB. **C-D)** Log2-assay expression of the *SIT1* (CR) and *RPC10* (TFIID) core promoters comparing genomic control sequences to UAS sequences, as in Figure 1C-D. **E)** Plot of log2-assay expression vs. expression rank for the four tested core promoters paired with all UAS sequences. **F)** Reporter expression (RT-qPCR of uGFP normalized to exogenous *S. Pombe* spike-in, using signal for the *TUB1* transcript) for nascent uGFP reporter controlled by four core promoters with and without genomic UAS. Fold induction is shown, calculated as the mean expression of core with UAS divided by mean expression of the core alone.

**Fig S2.**
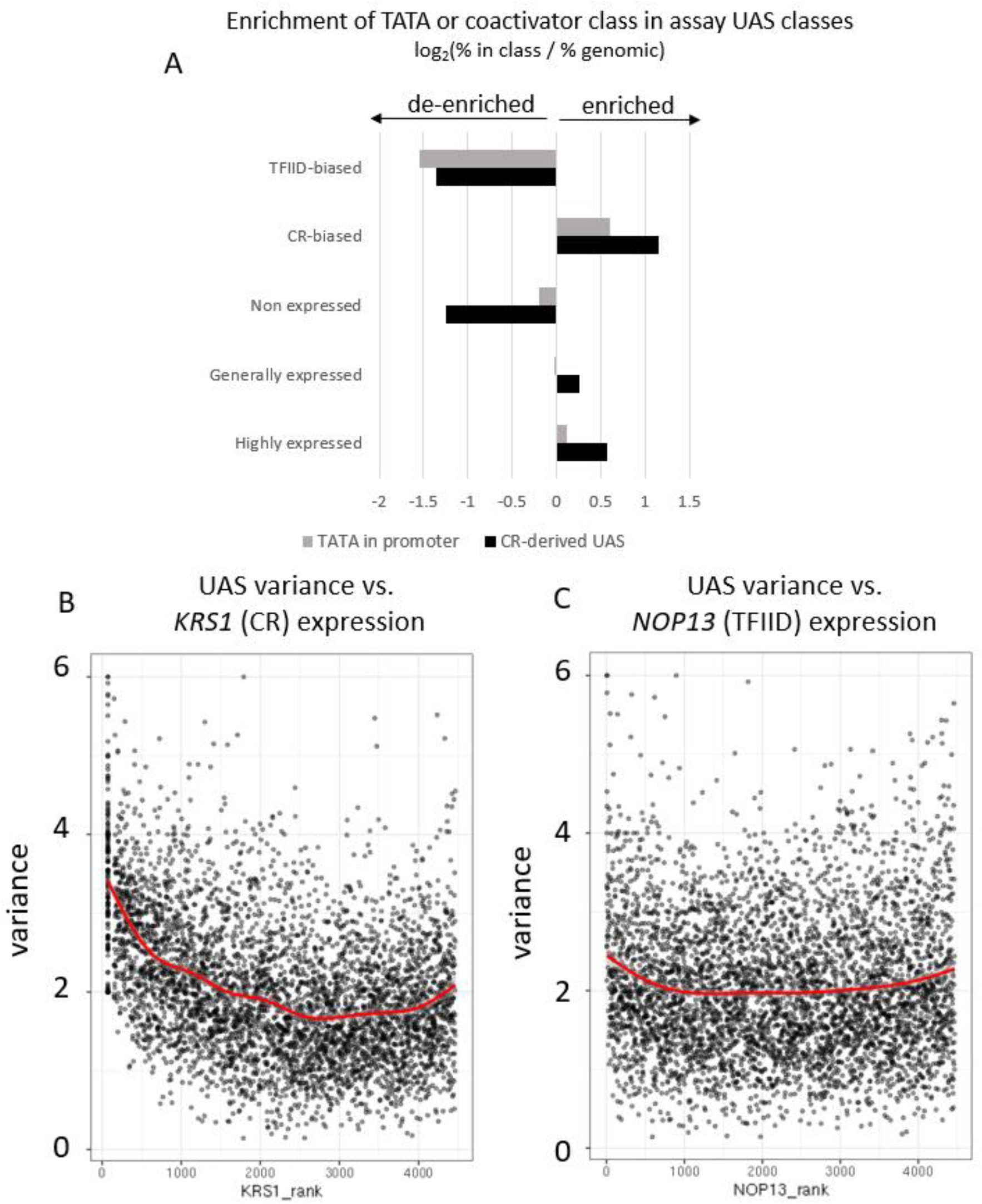
Log2-enrichment in percentage of UASs from CR-class genes or TATA-containing genes in UAS-core assay defined classes, compared to genomic percentage of CR class/TATA-containing genes. **B-C**. Plot of calculated UAS variance (see Methods) for all UASs vs. rank expression for UASs paired with *KRS1* or *NOP13* core promoter.

**Fig S3.**
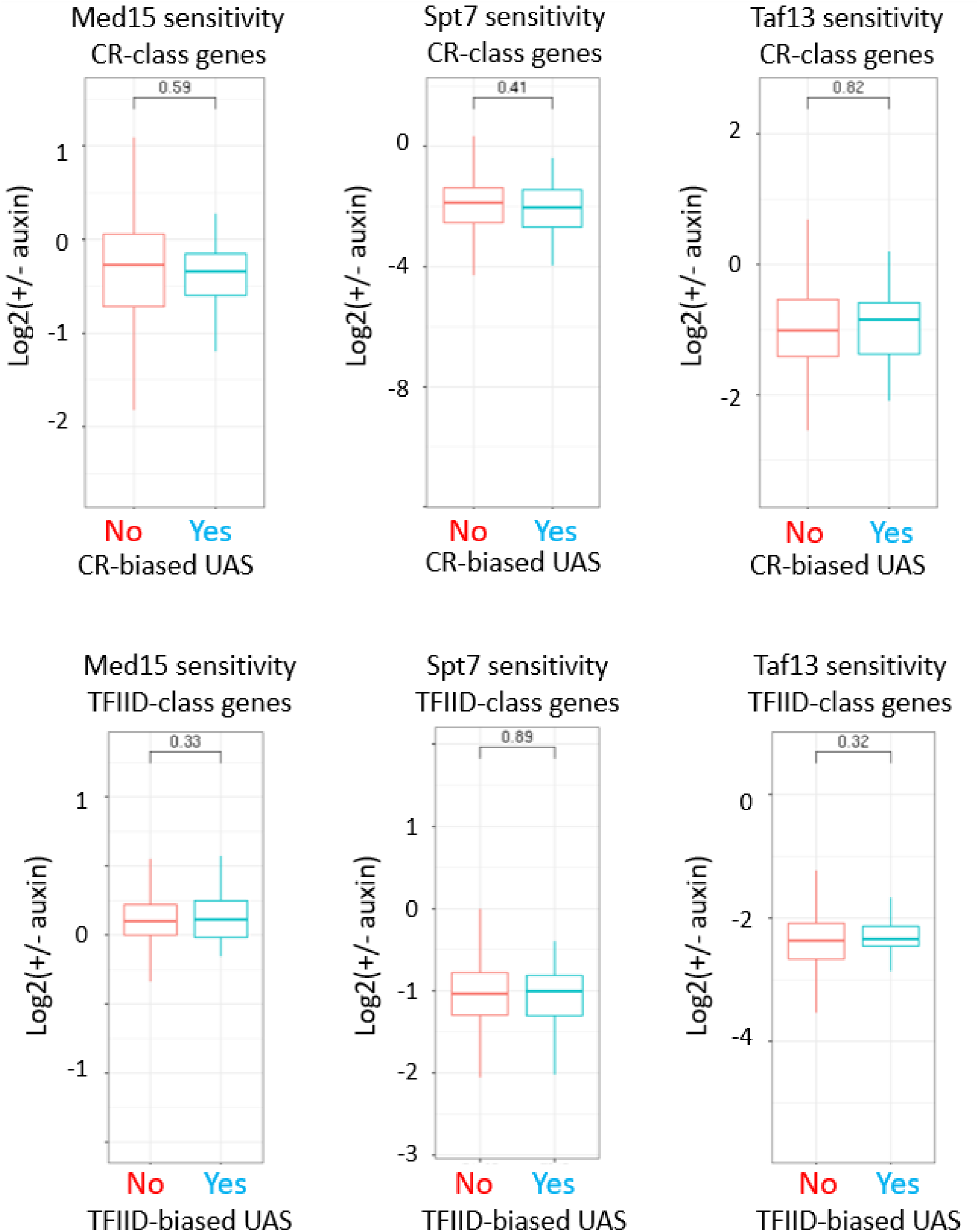
Transcriptional sensitivity to rapid coactivator depletion (MED tail/Med15, SAGA/Spt7, TFIID/Taf13) for endogenous genes, comparing UASs identified as CR-biased or TFIID-biased in UAS-core assay. Log2-FC after rapid depletion of coactivator subunits is shown for CR-class genes (top row) or TFIID-class genes (bottom row) (Donczew 2020, Warfield 2022).

**Fig S4:**
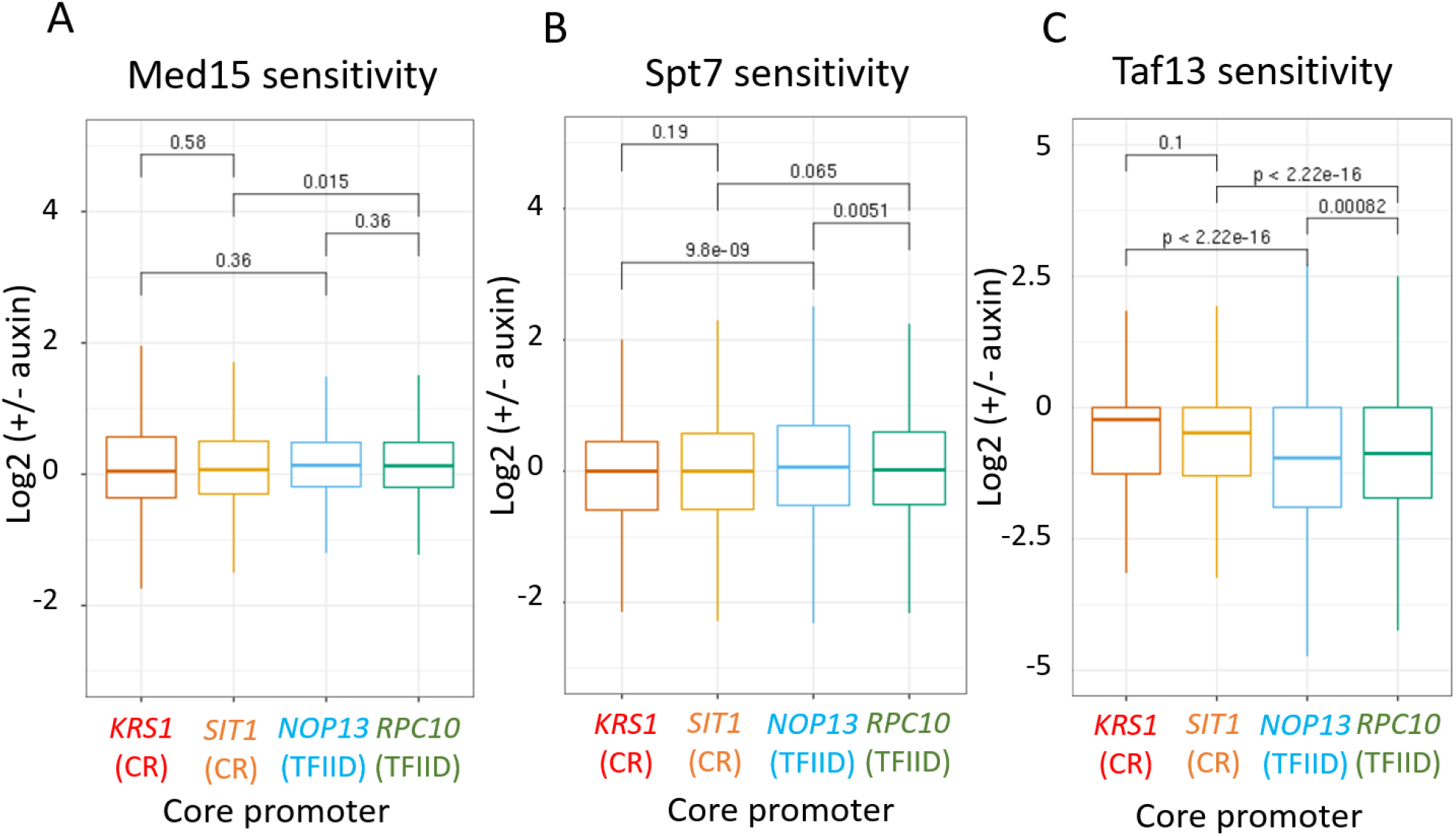
Coactivator sensitivity dependence and core promoter classes. **A)** Log2-fold change in transcription for all UASs paired with the four test promoters after Med15 depletion. T-test statistic is shown. **B)** Log2-fold change in transcription after Spt7 depletion. **C)** Log2-fold change in transcription after Taf13 depletion.

